# WlzWarp: An Open-Source Tool for Complex Alignment of Spatial Data

**DOI:** 10.1101/2022.02.11.480105

**Authors:** Bill Hill, Zsolt Husz, Chris Armit, Duncan R. Davidson, Paula Murphy, Richard A. Baldock

**Affiliations:** MRC Human Genetics Unit, Institute of Cancer and Genetics, University of Edinburgh, 4 Crewe Road EH4 2XU Edinburgh, UK; School of Natural Science, Trinity College, College Green, Dublin 2, Ireland; Deblur Ltd., Office 2, 30/2 Eskbank Office Complex, Hardengreen Industrial Estate, EH22 3NX Dalkeith, UK; BGI Hong Kong, 26/F, Kings Wing Plaza 2, 1 On Kwan Street, Shek Mun, NT, Hong kong

**Keywords:** 3D Images, Image Registration, Spatial Mapping, Gene Expression, Mouse Embryo, Non-linear Transforms

## Abstract

**Background:** The spatio-temporal organisation of many biological processes such as gene-expression and neuronal activity is critical to understanding the overall biological behaviour, phenotype and disease. This is especially true during embryonic development. During development spatial patterns of gene expression are key to segmentation, tissue differentiation and organ development. *In situ* techniques can reveal the activity of genes and the presence of proteins to a very high sub-cellular resolution but typically at high resolution only a few probes can be used on any sample therefore to compare many such patterns requires mapping of image-based data to a standard spatial framework. Once mapped those data can be collated, queried and analysed in purely spatial terms to reveal unknown combinatorial gene-activity that could not be discovered any other way. Mapping spatial data to image domains with systematic variation in shape and pose, such as embryos or elongated organs, presents special problems. Automated techniques available for more constrained systems can not deliver the mapping fidelity required to analyse these data therefore we have developed a manual editing tool, WlzWarp, for mapping 3D image data using the *constrained distance transform* (CDT) which uniquely can deliver the complex transforms required.

**Results:** We have implemented a fully open-source tool (available on GitHub), **WlzWarp** to provide interactive complex spatial mapping of 3D image data. We have applied **WlzWarp** to map a set of gene-expression patterns in the developing mouse embryo that could not be mapped by any other technique. The transform procedure was tested by using multiple images of the same gene from independent samples and thereby testing the full end-to-end mapping process. **WlzWarp** is implemented in C++ using open-source packages Qt, Coin and SIMVoleon for cross-architecture compatibility. It has been developed under Linux but also tested on Mac OSX and MS Windows.

**Conclusions:** **WlzWarp** is a freely available software tool for non-linear registration and alignment of complex 2D & 3D spatial patterns from one image to another or to an atlas model. It has been tested in the context of embryo data but can be used for any 3D or 2D image registration problem.

## Background

Enabling biological data to be collated, queried and analysed requires a common framework to provide a scaffold to align data from multiple observations and modalities. The genome provides the framework for varieties of ’omic data and ontologies have delivered annotation frameworks for aspects of biological location and function. Image-based spatial frameworks or atlases can capture spatially organised data such as gene-expression and phenotype at a higher location resolution than can be usefully described using ontologies. The use of explicit spatial atlases for gene-expression data was pioneered by the Edinburgh Mouse Atlas [1, 2, 3] and the development as an analytical framework described by [4]. The Mouse Atlas gene-expression database (EMAGE) [5] now holds over 30K curated and mapped gene-expression patterns across multiple mouse embryo developmental stages.

Many “atlas” based resources have been developed based on annotated images, here we focus on image data mapped onto a standard spatial atlas model to enable direct comparison of gene expression in terms of location and spatial patterning using image analysis techniques. If spatial data is mapped onto a common coordinate framework then direct spatial analysis can provide an objective approach to understanding gene-expression patterns. The Allen Brain Atlas has captured gene-expression data for virtually all genes expressed in the adult murine brain and a series of papers have shown the power of direct analysis independent of anatomical annotation [6]. These aspects and other analysis options including automated annotation and morphological variation are discussed by [7].

Spatial mapping from the original image to the coordinate framework is the critical step. For embryo data a complex non-linear mapping from 2D- or 3D-image data to the 3D framework is required. In many cases the transformation can not be undertaken with any of the tools developed for clinical and other biological data[8]. Furthermore these techniques do not admit manual override and editorial correction, and can not provide any form of real-time response or visualisation. For these reasons we developed a spline-based technique using the constrained distance transform (CDT) [9], which can deliver the spatial transforms required with a near real-time response to editorial changes. The CDT is the underlying algorithm for the developed WlzWarp application that provides a user interface for manual 3D image registration using the CDT.

The transformation technique CDT was compared with other algorithms in [9]. We do not know of any user-interfaces that can deliver real-time editorial control over 3D non-linear spatial mapping so direct comparison with other tools is not possible. We do however test the end-to-end accuracy and reliability by mapping multiple independent biological samples of embryos at a specific stage to show the well-understood gene-expression pattern of the gene *sonic hedgehog (Shh)*.

## Implementation

### Architecture

WlzWarp is a software application developed in C++ on a Unix architecture but with software libraries for the user-interface and visualisation aspects that allow re-compilation for both Mac OSX and MS-Windows operating systems. The core image manipulation and spatial mapping is provided by the Woolz[10, 11] image processing library. The user interface elements are provided using the Qt[12] C++ library which provides interface “widgets” with cross-platform support so that any application developed using Qt can be compiled to run under Linux, Mac OSX, MS Windows and Android with very little code change. The Image and volumetric model visualisation is provided by the Coin[13] C++ library with the SIMVoleon[14] extensions to deliver the required volume rendering. The WlzWarp application can be compiled to run under any operating system and hardware supported by Qt and Coin.

### Woolz

Woolz[10] is at core an image processing library developed in C. It includes a large set of command line tools for manipulating images and extensions for external data types and complex operations. It is fully open-source and available from GitHub[11] with documentation[15] provided using Doxygen. Woolz uses a unique image data representation combining an image *domain* encoding the spatial extent of the image, which can be an arbitrary point-set in 2D pixel or 3D voxel space, and a separate *values* table capturing the data values at each location within the domain. The interval-based domain representation has the advantage compared to other image encodings of a compact non-compressed in-memory encoding and very fast binary operations on the image data. These include the set operations of union, intersection and difference, morphological operation of dilation and erosion with arbitrary structuring elements.

Mesh domains are a 2D (triangles) or 3D (tetrahedra) representation of a domain conforming to the volume of space occupied by that domain and are used provide non-linear invertible spatial transforms[9]. A transform is represented as a mesh domain with values at each node that are vector displacement values. The full voxel transform is calculated by interpolation within the mesh primitives and can be computed fast enough to deliver interactive update of a full volume transform. The transform can encode any non-linear deformation including mappings that do no preserve topology.

### The Constrained Distance Transform

To deliver a rapid volumetric transform we use the mesh-based approach in combination with Radial Basis Function (RBF) transforms. RBFs are frequently used for interactive image warping and perform well for small deformations. However, when the deformation gradients are large these methods may produce non-diffeomorphic, mirrored or extremely distorted mappings[16]. Our method uses RBFs, but with geodesic rather than Euclidean distances. We call this the constrained distance transform (CDT) which has been fully described by Hill and Baldock (2015)[9] and delivers the complex spatial mapping required for spatial data mapping in the embryo coupled with the interactive transform capability afforded by using a mesh-based approach.

### Methodology

WlzWarp provides an interactive graphical user-interface (GUI) to spatially map via a non-linear and non-diffeomorphic transform a source image, *Is*, to a target image, *It*. The images can be either 2D or 3D and in the atlas application the target is typically the model used to capture the spatial patterns from many observations as in the Emage[5] database for *in situ* gene-expression data. The interface (see fig 1) provides three windows with 3D visualisation of: the source image; the target image and the merged image with either the source mapped onto the target (forward transform) or the target mapped onto the source image (reverse transform). Within each window the user can control the visualisation in terms of colour mapping and transparency, shaded surfaces or wireframe, volumetric rendering with graded and bounded opacity.

**Figure 1.**
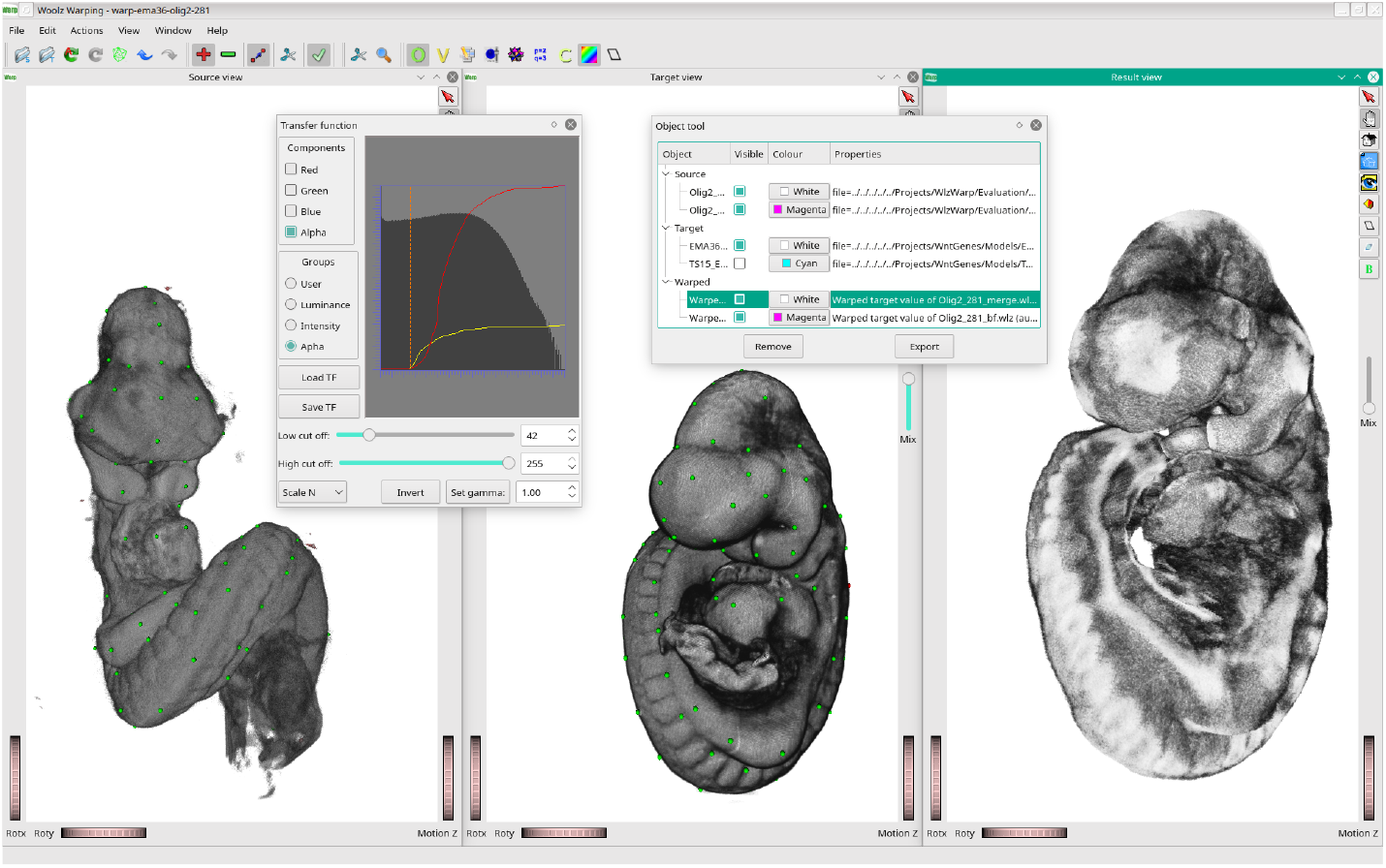
The WlzWarp graphical user interface. The left-hand window shows the source 3D image rendered to show the outer surface of the volume. The centre window shows the target volume rendered in the same way. The green dots show the position of the tie-points entered manually pair-wise to define source-target correspondence. The right-hand window shows the source image transformed to the shape and pose of the target. In this example the left-handed twist of the source has been transform to the right-handed twist of the target which is a transform not possible to achieve with other warping techniques.

The transformation utilises the RBF in the context of a conforming mesh defined usually on the target image and which is typically pre-calculated and optimised. The user then defines the required transform by placing a series of “tie-points” or “landmarks” which are points on the source and target that correspond i.e. are matching locations. On placement of a landmark pair the transformed object is updated and displayed either automatically or on demand according to a user-setting. Landmarks are added sequentially and adjusted until a correct registration is achieved. This is a purely manual registration process designed to provide interactive control to the domain expert. Landmarks can be placed at any pixel or voxel location either directly within the visualised volume, a virtual section within the volume or to a defined surface. For surface placement a “snap to surface” function provides an efficient mechanism to ensure the point is correctly placed on the surface of the object.

The selection of tie-points will depend on the application. For embryo data with large conformational differences we have found a staged approach to be most productive. In the first stage approximately 10-20 tie-point pairs are identified using unambiguously identifiable anatomical points for example the anterior tip of the notochord (when still visible), a position on the dorsal midline of the neural tube at a specific somite number or locations of the optic and otic vesicles. This first stage provides a primary alignment and followed by tie-points associated typically with surface features such as tips of the limb-buds or the branchial arches then finally the surface and volume alignments can be adjusted using tie-points selected proportionately between other points and guided by the quality of the visualised mapping.

To allow image registration a conforming mesh is used. This is either a triangular mesh for 2D or tetrahedral mesh for 3D and the mesh boundary must conform to the boundary of either the segmented target or source image domain. The purpose of this mesh is two-fold; to allow rapid evaluation of the RBF and to compute distances along a path constrained to the domain. For 2D and very simple 3D images the mesh may be generated within WlzWarp but for more typical 3D images an external mesh generator such as netgen[17] is required.

Each session of image registration is tracked and saved via a project file and directory structure which allows recording, re-starting and sharing of the registration detail. WlzWarp provides export functions for the mesh transform which can be used to transform any woolz object with a domain within the domain of the transform mesh. These transfomations can also be applied outside of the WlzWarp application and forms a core operation for automated mapping of spatial data for the Mouse Atlas gene-expression database.

## Results

Here we present examples of the use of the WlzWarp interface for mapping of both 2-D and 3-D image data selected to illustrate how the CDT can deliver the required mapping and a comparison with the outcome not using the CDT.

### 2D Warp of Embryonic Wholemount Images

This exemplar test of the CDT illustrates the capability to disambiguate overlapped data in a 2D image onto a target image using a transform that does not preserve topology. A wholemount assay image was selected from the EMAGE gene-expression database (showing HOXB6 expression[18] see figure 2) along with the appropriate E9.5 (Theiler stage 15) atlas model 2D view. In EMAGE the TS15 2D wholemount views have the tail articulated to allow independent spatial mapping to the embryo tail and the remainder of the body. A version of the wholemount target image was created in which the object domain was no longer segmented from the background. This was done by simply making a rectangular object enclosing the model image and using that rectangular mesh to calculate the transform. With the assay image as source image and the appropriate wholemount view as target image two projects were created using WlzWarp. The same basis function parameters were used in both cases. Other than the target images, the only difference between the projects was in the automatically generated conforming meshes. Because the mesh generated for the rectangular domain did not conform to the required target domain, distances used to evaluate the basis function mesh displacements were not constrained to the target domain. Views of WlzWarp showing the source and target images, the target meshes and the transformed source images are shown in figure 2. These show that the CDT will segment a source image (using the already segmented target mesh) and may produce one-to-many mappings where appropriate(fig. 2c). The resulting transformation when distances were no longer constrained to the target domain is neither segmented from it’s background nor well registered to the target image (fig. 2d).

**Figure 2.**
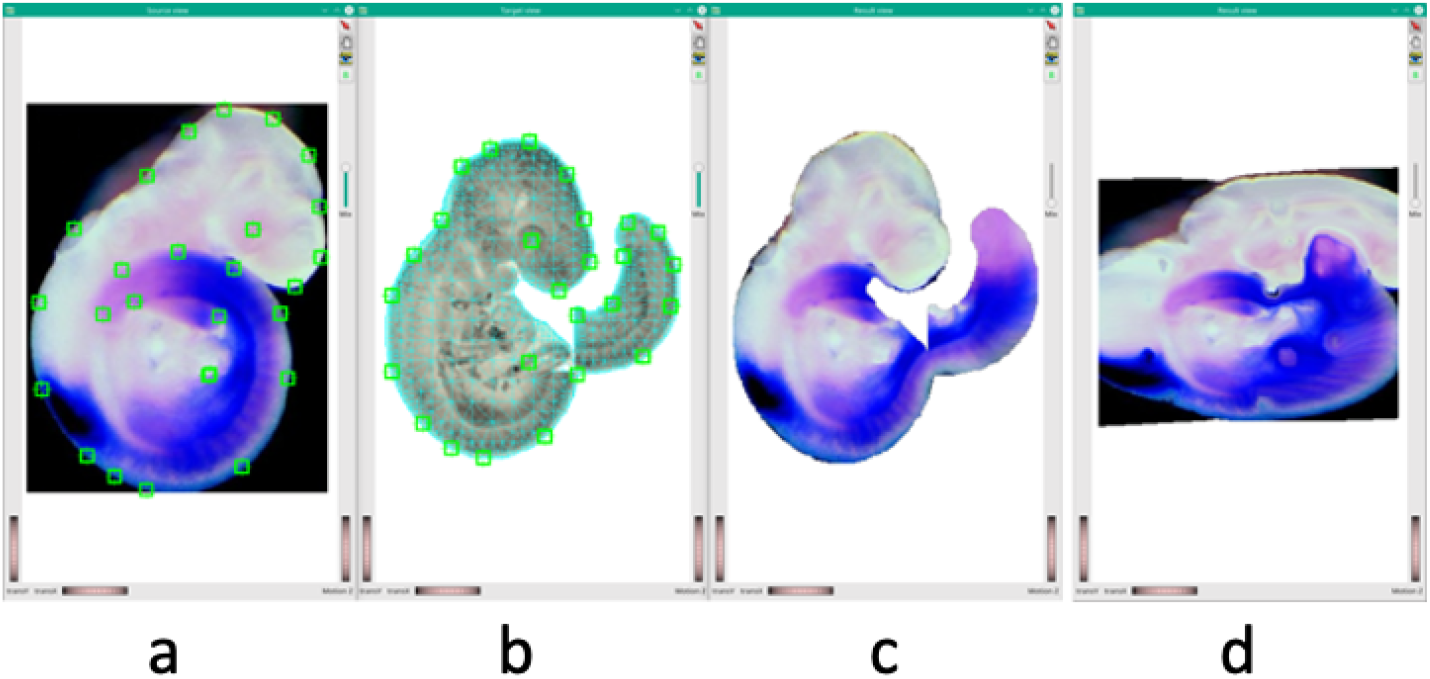
Screenshots of the 2D registration process using WlzWarp for both conforming and non conforming meshes. (a) shows the original expression image with the selected tie-point locations shown as small squares, (b) shows the target model image with offset “tail” and the matching tie-point correspondences. The boundary of the constrained mesh is shown as a green line around the edge of the embryo view. (c) is the embryo mapped with the conforming mesh and with radial basis function transform displacements using geodesic distances within the mesh. (d) Traditionally mapped embryo using a non-conforming mesh in which the distances are Euclidean within the plane and not constrained to be within the embryo.

### 3D Warp of Embryonic Spatial Data

The purpose of this exemplar is to show how the CDT can transform a set of 3D volume images of a biological entities that present with radically different conformations sufficient to defeat other techniques developed for this type of volumetric registration[9]. A set of nine paired optical projection tomography images of Theiler stage 15 (TS15) mouse embryos, with each embryo being stained to show gene expression, was selected to form an evaluation dataset, with the image pairs consisting of autofluorescence and brightfield images. The brightfield image is captured to show the regions of expression and the autoflourescence image reveals the embryo anatomy. Within the autoflourescence image the expression region shows”drop-out” of signal so the images are merged to generate a complete volume for the registration process. For the purposes of registration the tie-points are all associated with anatomical landmarks and surface locations and none use the expression pattern. Figure 3 shows 3D rendering of the merged images. The top-row of embryos in figure 3 have also been rendered to show the expression pattern (brightfield image) in the context of the whole embryo (autoflourescent image).

**Figure 3.**
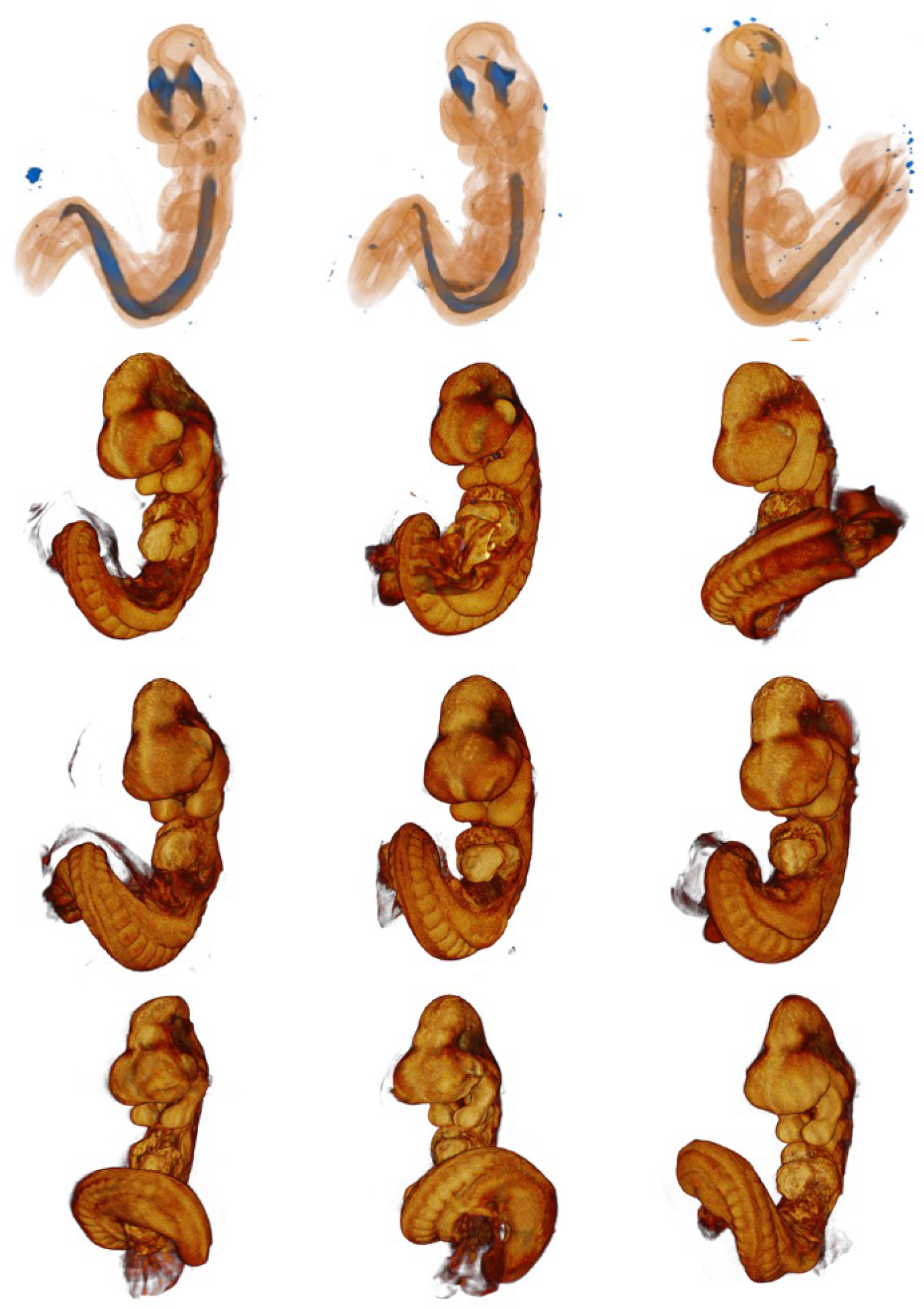
3D embryo dataset in which each embryo shows the combined reference and signal components. These are a set of embryos from two litters and show the typical range of pose, presentation and developmental variation typical of the E9.5 embryos. The top row of images is rendered with transparency to show the extent of the *Shh* expression pattern within each of the embryos depicted in the second row. Rows three and four have a similar internal shh expression.

The dataset was selected from the same developmental stage to show the wide varieties of pose and presentation typical of this stage with some additional developmental variation. The autofluorescence images were merged with the brightfield images to form a reference image for each embryo. It is very clear in figure 3 that the trunk and tail can present either as part of a right-hand or a left-hand spiral. We have shown earlier [9] that this transformation defeats any system not-based on a conforming mesh. The EMAGE TS15 OPT atlas model (EMA36) and it’s associated conforming mesh were the target for the evaluation embryo set. Using WlzWarp for all nine assay images landmark pairs were placed on the merged source images and the atlas target image. The overwhelming majority of these landmarks were placed on the embryo surface with few being placed internally within the 3D image volume. Table shows the number of tie-point placements required for each of the nine-embryos used for this analysis to provide an idea of the effort required to map each embryo. Typical user-time for mapping each embryo was in the range 30-60 minutes.

**Table 1.** Table of the numbers of tie-points used for mapping the embryos shown in figure 4. Excluding the embryos showing expression the embryos are numbers sequentially from the top-left (embryo 1) across the rows to the bottom-right (embryo 9). The values of the inverse multiquadric control (IMQ) parameter delta (see [9]) are provided for each embryo in the third column as examples for using WlzWarp.

| Embryo | Tie-points | IMQ Delta |
| --- | --- | --- |
| 1 | 80 | 0.04 |
| 2 | 82 | 0.04 |
| 3 | 92 | 0.04 |
| 4 | 78 | 0.04 |
| 5 | 83 | 0.04 |
| 6 | 85 | 0.04 |
| 7 | 98 | 0.08 |
| 8 | 138 | 0.075 |
| 9 | 101 | 0.075 |

Figure 4 shows the warped 3D embryo dataset of figure 3 after registration.

**Figure 4.**
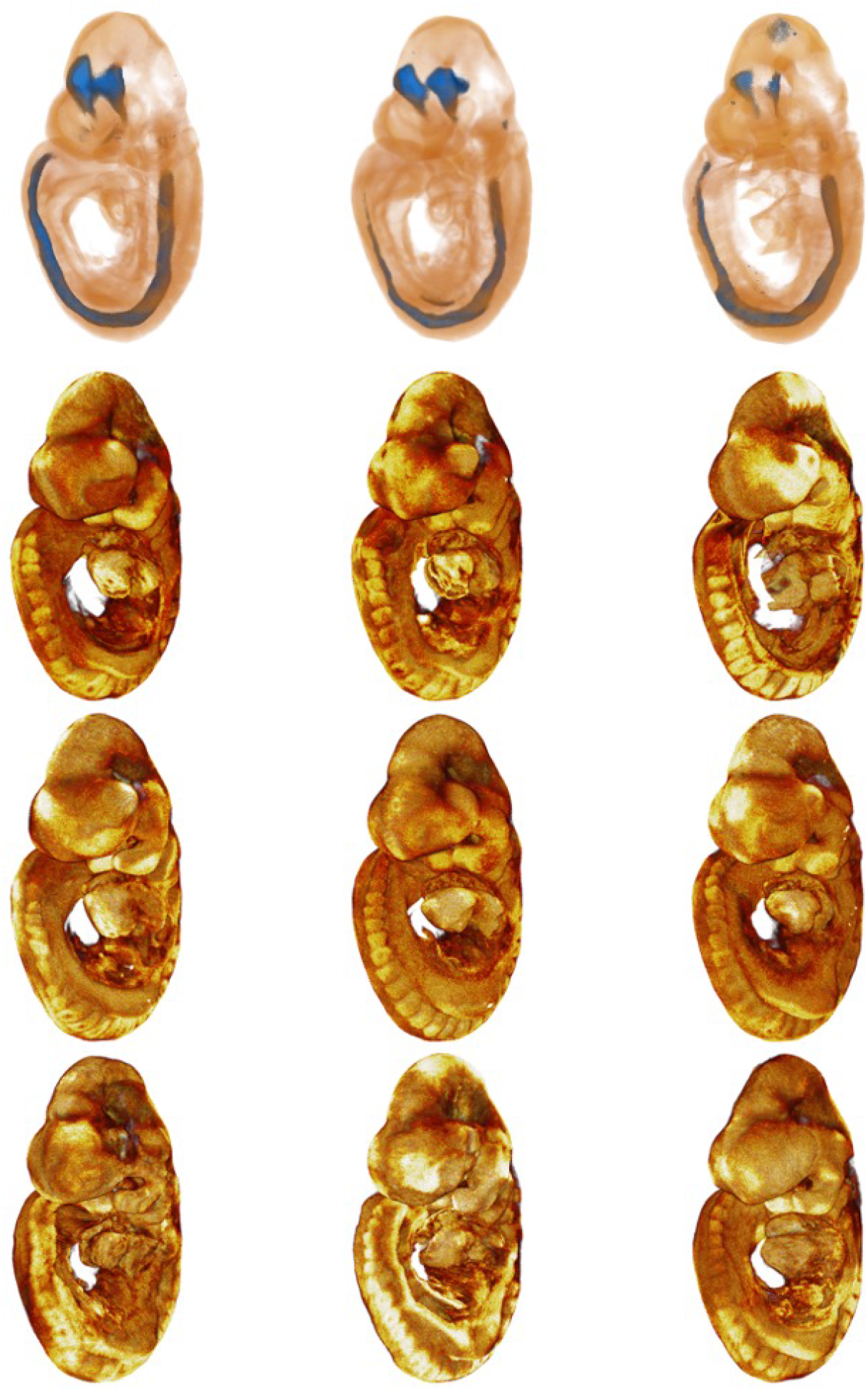
Results of using WlzWarp to map each of the embryos shown in figure 4 to the EMAGE 3D model for the E9.5 developmental stage. All pose variations are correctly mapped. The top row of embryos show the mapping of the expression domains for each of the embryos in the second row respectively.

To illustrate the quality of the non-linear registration process using WlzWarp we use the very consistent and repeatable pattern of expression of the gene. Each brightfield image visualises the expression pattern but with variation in actual intensity arising from variation in the histological and imaging processes. To extract the expression each brightfield image was histogram matched to one of the embryos then thresholded using a single fixed threshold value. This results in a spatial domain enclosing the region of signal for each embryo. This spatial domain was then transformed using the mapping established by WlzWarp independently for each of the nine embryos. The subsequent domain images were then merged to form an “occupancy” image by calculating for each voxel in the 3-D volume the number of times that location was within the nine mapped expression patterns. This occupancy image is displayed as a volume rendered image in figure 5 with selected sectional cuts through the volume to show the spread of occupancy values.

**Figure 5.**
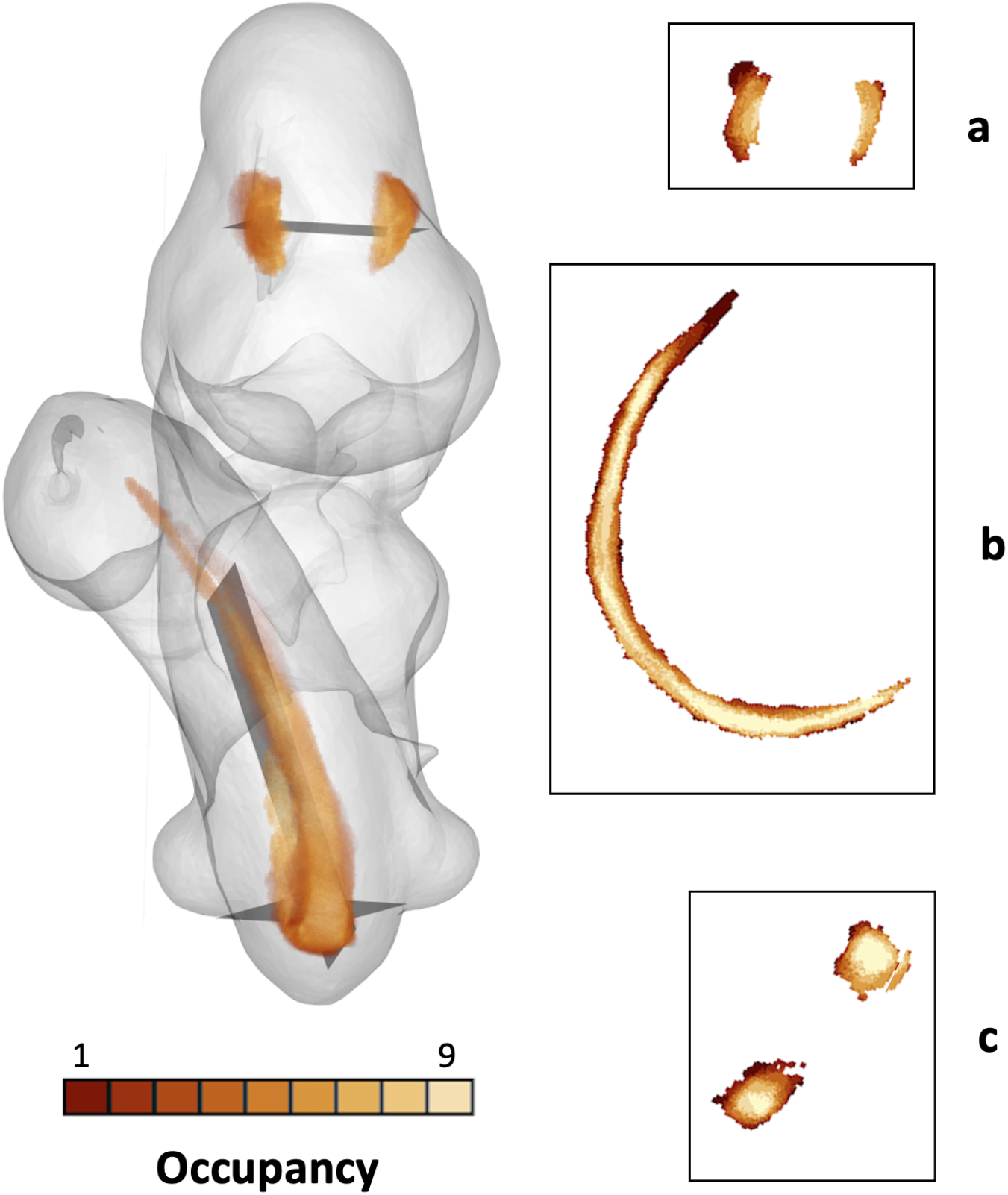
Expression occupancy of nine combined *Shh* expression domains. The overall 3D occupancy is shown on the left with selected sections through the 3D occupancy data shown on the right. Occupancy values within the sections are indicated using the heat-map colour scale. **(a)** shows a transverse section through the head in the region of the diencephalon; **(b)** a longitudinal/oblique section though the neural showing occupancy from the vicinity of the forelimb-buds through to the tail region; **(c)** a transverse section through the neural tube close to the embryo rump.

Given the tightly localised and repeatable nature of ssh expression patterns, the spread of occupancy values is therefore a good indication of the mapping capability of the CDT. The section views in figure 5 show patterns that are high (6+) occupancy to within a few voxels of the pattern edge suggesting a mapping error of the pattern boundaries of less than 5 voxels. This shows very high consistency: more remarkable because none of the mapping used the expression pattern for alignment, i.e. all mapping correspondences (tie-points) were identified using anatomical features.

## Discussion

The constrained distance transform (CDT)[9] uses a radial-basis function in combination with a mesh-based tessellation of a 2D and 3D space and geodesic distances calculated within the constraints of the spatial domain of interest. This delivers an interactive warping capability enabling complex and non-diffeomorphic (non-topology preserving) mapping of 2D and 3D image data. Here we have presented a graphical user-interface, *WlzWarp*, that utilises the CDT to provide a convenient manual editing tool for spatial data-mapping and illustrate its use in the context of biomedical spatial data. It can of course be used for any application requiring complex non-linear mapping of spatially organised data. All of the software developed for WlzWarp is fully and freely available open-source from the MATech github repository.

The results reported here illustrate the capabilities of the warping technique and the utility of the WlzWarp GUI and the typical level of accuracy the tool can deliver when applied to a set of mouse embryo gene expression data for the *Shh* gene. The level of accuracy that can be achieved is primarily dependent on the number of tie-point correspondences entered and the knowledge and expertise of the user. WlzWarp is most useful with image data for which automated methods fail including *in situ* gene-expression data as the expression signal can obscure underlying structures and provides conflicting patterns that would confuse any pattern matching algorithm.

WlzWarp has been used for a large number of studies requiring complex non-linear mapping of spatial data. All of the 3D mapped for the EMAGE [5] database has been registered against the model embryos using the CDT primarily using the WlzWarp application making possible the spatial query and similarity search and analysis. A study of chick heart development[19] based on clusters of highly expressed genes provides a comprehensive structure-based view of genes showing significant expression levels. The “straight mouse embryo” [20] illustrated the mapping complexity that can be achieved and how “natural” biological coordinates relating to the underlying biological structures can be mapped onto the 3D cartesian frame of the embryo in “real” space. WlzWarp was also critical or an extensive analysis of Wnt signalling pathway genes enabling a unique embryo-wide spatial analysis across developmental stages, revealing complex co-expression patterns among Wnt, Fzd and other pathway gene family members (submitted by these authors for publication).

## Conclusions

The constrained-distance transform provides a unique mapping capability not delivered by other tools for mapping spatial data. WlzWarp is an application harnessing the CDT with a customisable GUI to allow mapping of data with real-time editing of the transformation and is particularly useful for data that resists other automated and manual systems. In particular this allow mapping of valuable data that would otherwise be lost by not fitting the tight presentation and pose constraints required for other systems.

WlzWarp and all underlying Woolz software is full available from GitHub with details of third party software dependencies and how to compile the application from source. It is a complex application with many options for controlling the visualisation and output. There are some help files available and further questions can answered by contacting the corresponding author.

## Acknowledgements

The authors thank The Company of Biologists for permission to use figure 3D from Akasaka *et al* [18] licence ID: 1169681-1.

## Abbreviations

RBF: radial basis function
GUI: graphical user-interface

## Availability and requirements

**Project name:** WlzWarp

**Project home page:** github.com/ma-tech/WlzQtApps

**Operating system(s):** Linux, Mac OSX, Windows

**Programming language:** C++ & C

**Other requirements:** open-source packages SIMVoleon, Coin and Qt

**License:** GNU GPL 2

**Restrictions:** None

## Authors’ contributions

BH - conceptual development, software design, development and implementation and of the CDT technique, execution of the mapping experiments and manuscript preparation; ZH - software development and interface implementation; CA - trials and testing of WlzWarp and data mapping and manuscript preparation; DRD & PM - critical review for embryo mapping, provision of image data and manuscript preparation; RAB - software and conceptual development, manuscript preparation.

